# Anti-fibrotic activity of an antimicrobial peptide in a *Drosophila* model

**DOI:** 10.1101/2021.03.19.436168

**Authors:** Dilan Khalili, Christina Kalcher, Stefan Baumgartner, Ulrich Theopold

**Author notes:** Corresponding author: Ulrich Theopold, Department of Molecular Biosciences, The Wenner-Gren Institute (MBW), Stockholm University, Svante Arrheniusväg 20c, 10691 Stockholm, Sweden.

## Abstract

Fibrotic lesions accompany several pathological conditions including tumors. We show that expression of a dominant-active form of the Ras oncogene in *Drosophila* salivary glands (SGs) leads to redistribution of components of the basement membrane (BM) and fibrotic lesions. Similar to several types of mammalian fibrosis, the disturbed BM attracts clot components including insect transglutaminase and phenoloxidase. SG epithelial cells show reduced apico-basal polarity accompanied by a loss of secretory activity. Both the fibrotic lesions and the reduced cell polarity are alleviated by ectopic expression of the antimicrobial peptide Drosomycin (Drs), which also restores secretory activity of the SGs. In addition to ECM components, both Drs and F-actin localize to fibrotic lesions.

## Introduction

Extracellular matrices (ECMs) are highly specialized three dimensional, tissue- and stage-specific structures, which surround cells. Much of our understanding of ECM structure and dynamics has been obtained by studying model organisms [1, 2]. The ECM is composed of fibrous as well as non-fibrous proteins and of proteoglycans. Fibrous collagen and laminin form two extracellular networks in part through self-assembly [3, 4]. Non-fibrous proteins of the ECM including nidogen and the proteoglycan perlecan mediate the interaction between the collagen- and laminin networks [4]. Formation of the ECM is influenced by the underlying cells via receptors such as dystroglycan, syndecans and integrins, which mediate contact between the cells and the ECM [3, 5, 4]. Mutual cell-ECM interactions influence cell adhesion, migration, proliferation, apoptosis and cell differentiation during development. During recent years, it has been increasingly recognized that instead of being a stable structure, the ECM is a highly dynamic matrix, which undergoes constant turnover both during development and as part of physiologic adaptations [3, 5, 4]. During development, the ECM directs morphogenetic processes for example in the intestine, the lungs and the mammary and submandibular (salivary) glands [3, 5]. Physiological responses include rapid repair upon tissue damage, which restores cellular and ECM integrity. In contrast, if unchecked, ECM remodeling in humans may promote pathological states such as osteoarthritis, fibrosis and progression towards cancer [3, 6, 7, 5] and may impede the delivery of drugs [8]. Fibrosis affects different organs including the skin, liver, kidney, heart and the lungs [9]. Fibrotic triggers include genetic disorders, mechanical stimuli, such as asbestos, poorly controlled diabetes and hypertension, tissue dysplasia such as in tumors as well as persistent infections [10]. Pulmonary fibrosis may be of unknown origin (idiopathic, [11]) or a consequence of infections such as during severe cases of Covid-19 [12]. Fibrotic lesions are often characterized by an unabated activation of wound healing [10]. Chronic activation may occur during the (1) initial hemostatic phase, the subsequent (2) recruitment of cells of innate and adaptive immunity or (3) during the remodeling and restoration of tissue integrity. The last stage includes the migration and differentiation of fibroblasts into myofibroblasts [9]. Fibroblastic pathologies include increased production of ECM components, increased stiffness of the ECM as well as chronic immune activation and inflammation [9, 3, 8, 4]. ECM dysregulation often affects tissue function and may lead to complete organ failure and ultimately to death [10, 13, 14, 4].

Effector mechanisms of insect innate immune systems comprise both soluble and cellular reactions [15, 16]. Soluble mediators are released from immune-competent organs, primarily the fat body but also the gut, the tracheae and the salivary glands [15, 17-19]. Among the most strongly induced soluble mediators are several classes of antimicrobial peptides (AMPs), with both antibacterial and antifungal activity [15]. One example for the latter is the antifungal peptide Drosomycin (Drs, [20]). Cellular immunity relies on hemocytes, immune cells in the hemolymph, which in non-infected *Drosophila melanogaster* comprise plasmatocytes and crystal cells [15, 21, 22]. Plasmatocytes have been likened to macrophages although recent single cell RNAsequencing revealed a high level of diversity among plasmatocytes [16, 22-24]. They are also key players during insect hemostasis and release coagulation factors, which help to seal wounds and prevent dissemination of microbial intruders [25]. Coagulation factors in insects include transglutaminase, a homologue of the mammalian clotting factor XIIIa [26, 27, 22]. Crystal cells contain inactive precursors of the enzyme phenoloxidase (prophenoloxidase, PPO), which is activated upon infection and wounding and leads to the production of intermediates with antibacterial and crosslinking activity at wound sites, and ultimately to the production of melanin [22]. Melanization is also the final part of responses against large numbers of bacteria (nodule formation) and against larger intruders such as wasp eggs (encapsulation, [15, 16]). Nodule formation and encapsulation have been likened to the formation of granulomas in vertebrates [28, 29, 25]. Melanization is also observed as a consequence of deregulation of immunity and as part of the response against aberrant self-tissues similar to autoimmune reactions in mammals. Mutants that display endogenously-driven melanization have been named “melanotic tumor mutants” although the underlying mechanism is not necessarily linked to mutations in tumor-promoting genes [30].

Dysregulation of the ECM with similarities to fibrosis has been induced experimentally in *Drosophila* and the effects on immunity studied [31]. In flies, one particular kind of ECM, the basement membrane (BM) covers all internal organs and separates them from the hemolymph in the open circulatory system. The BM is created through the release of its components from the fat body into the hemolymph and subsequent deposition onto the basal site of tissues including the fat body itself. Hemocytes contribute to BM formation or its repair [32-37] although the relative contribution of hemocytes versus fat body may be variable [38]. In one study, the formation of BM deposits with similarity to fibrotic lesions was induced in the fat body by plasma membrane overgrowth or alternatively through increased secretion of immune effectors. Both scenarios led to a damage response including melanization [31]. Melanization was also observed by simultaneous disruption of the basement membrane and loss of cell polarity [39]. Similarly, the *Drosophila* ECM protein SPARC (Secreted Protein, Acidic and Rich in Cysteine, also called BM40) was found to contribute to age-related cardiac fibrotic deposits, associated with reduced life span in flies [40]. Here, we show that ECM components are dysregulated in flies that express a dominant-active oncogene (Ras^V12^) in salivary glands (SGs; [29]). SGs have so far been mostly studied as models for organ development [41-44] and for innate immunity [45]. Proper secretion of salivary glue proteins and other mucin-type secretions in preparation for pupation is indicative of normal SG function [44] and is expected to be negatively affected by the formation of fibrosis [4] and this is what we observe. The fibrotic phenotype is strongest in the distal, secretory part of the SGs where ECM dysregulation coincides with the recruitment of transglutaminase and phenoloxidase and of hemocytes. In contrast, the proximal part of SGs displays an almost normal histology despite strong activation of AMPs, most notably the AMP Drosomycin. Due to the similarities to fibrotic lesions in mammals we refer to the regions with dysregulated ECM deposition concomitant with inflammatory reactions [46] as fibrotic deposits, although we acknowledge some differences to mammalian fibrosis [47]. Surprisingly, forced SG-wide expression of Drs including distal SGs strongly reverted fibrosis and restored tissue integrity in distal SGs and secretory activity to almost normal levels. This indicates that depending on the type of response, innate immune reactions may have pro- as well as anti-fibrotic consequences and establishes AMPs as regulators of tissue homeostasis.

## Material and methods

### Fly strains and sample preparation

*w*^*1118*^,*Beadex*^*MS1096*^*-Gal4* (Referred here as *Bx*: 8860/Bl), *w*^*1118*^, *w*^*1118*^;;*UAS-Ras*^*V12*^ (4847/Bl), *w*^*1118*^;;*UAS-Drs* (*Drs-OE*: overexpression [46], *w*^*1118*^; *Vkg* ^*G00454*^ */CyO*.*GFP [48], UAS-Drs-HA* [49], *Bx,mCD8::RFP* [46] and *Sgs3-GFP* (5884 /Bl), (see also supplementary table S1). Flies were cultured in 25°C, 12 h dark/light cycle room. Female virgins were collected for five days, crossed to the respective male on day 7. Progeny larvae were grown as described in [46]. 10-20 salivary gland pairs were fixed in 4% PFA for 20 min. Samples subjected for extracellular staining were washed 3 x 10 min with 1x Phosphate buffered saline (PBS). Samples used to stain for intracellular expression were washed 3 x 10 min with PBS-T (PBS with Triton-X100: 1%) before staining.

### Production of antibodies

GM02366 (Flybase, [50]) coding for *Drosophila* SPARC was used as a template for PCR amplification using Takara Ex Taq polymerase (Takara Biomedicals) according to the manufacturer’s instructions. 5′ and 3′ primers equipped with suitable restriction enzymes created a PCR fragment spanning the complete open reading frame lacking the SPARC signal peptide that was subsequently ligated in-frame downstream of the mouse BM-40 signal peptide of the episomal expression vector pCEP-Pu [51]. After verification of the sequence, the expression vector was used to transfect human 293-EBNA cells (Invitrogen) and serum-free medium was collected for protein purification according to established methods [51]. Immunization of rabbits, affinity-purification of antibodies and ELISA titration followed standard protocols [52].

### Immunohistochemistry

Antibodies against SPARC (1:3000:this report), Nidogen (1:2000) [53], Laminin (1:2000) [54] and Perlecan (1:2000) [55] were incubated overnight (ON) at 4°C either in PBS (non-permeabilized) or PBS-T (permeabilized). PPO1 (1:500), PPO2 (1:250), Tiggrin (1:500, [56] a kind gift by A. Simmonds, Alberta) were stained in PBS at 4°C O/N. Anti-ß-integrin (1:200, DSHB #CF.6G11) and Anti-Dlg (1:50, DSHB #4F3, [44]) was incubated at 4°C, O/N in PBS-T. Anti-HA (1:3000, Thermofisher #5B1D10) was pre-incubated with fixed *Ras*^*V12*^ salivary glands for 1h at RT in PBS-T and subsequently incubated with the samples for 1h at RT. Samples were washed 3 x 10 min with PBS/PBS-T and incubated with secondary antibody anti-Rabbit-568 (1:500, Thermofisher #A21069), anti-Rabbit-488 (1:100, Thermofisher #A11008) or anti-Mouse-546 (1:500, Thermofisher #A11030), DAPI (1:500, Sigma-Aldrich D9542) and Phalloidin-488/546 (1:500, Sigma-Aldrich #A12379 and # A22283, respectively) for 1 h at room temperature RT, washed 3 x 10 min with PBS or PBS-T before mounting in FluoromountG.

### Transglutaminase activity

Salivary glands were fixed in 4% PFA for 20 min and washed 3 x 10 min with PBS at RT. The glands were incubated with TQ, an antibody that recognizes isopeptide (ε-(ϒ-L-glutamyl)-L-lysine) generated by transglutaminase (1:100, Covalab #mab0012), in PBS at 4°C O/N. Samples were washed 3 x 10 min in PBS. Primary antibody was detected with anti-Mouse-546 (1:500, Thermofisher #A11030) for 1 h at RT. Thereafter, the samples were washed 3 x 10 min in PBS and subsequently mounted in FluoromountG. Image acquisition and analysis is described below.

### Image acquisition and analysis

Whole SGs were photographed with a Zeiss AxioscopeII microscope and images were exported as TIF files. The intensity was measured of the whole gland with ImageJ (ver. 1.52n). For co-variation of the intensity, the signal (SPARC, TQ, PPO1, PPO2 or HA) of the individual salivary glands was plotted against Phalloidin intensity signal of the same gland. Representative images were captured with a confocal Zeiss LSM780 microscope. Final figures were made on Affinity Designer (version 1.7.3). Z-stack images were taken with LSM780 (objective 40x/1m3 Oil and 63x/1.40 Oil DiC M27) and co-localization analysis was analyzed on Imaris (version 1.52n, coloc module). Statistics was performed in GraphPad (8.3.0), including D’Agostino for normal distribution, unpaired t-test, Pearson’s correlation and linear regression.

#### Basal membrane thickness analysis

Z-stacked images were analyzed on a Y-Z plane in ImageJ (ver. 1.52n). The ‘Straight line’ tool was used to measure the thickness of Collagen IV (Vkg-GFP), presented in pixels (px). Ten measurements across a Y-Z plane were averaged per salivary gland. A minimum of tree salivary glands per genotype were measured. An unpaired t-test was performed in GraphPad (8.3.0).

### Reverse Transcription quantitative PCR (RT-qPCR)

RNA extraction, cDNA synthesis and qPCR analysis were performed as described in [46]. qPCR was performed on three independent replicas targeting *Rpl32, Sgs3* and *Eig73Ee*. Primer sequence for *Rpl32*: 5’-CGGATCGATATGCTA-3’ and 5’-CGACGCACTCTGTTG-3’, *Sgs3*: 5’-GTGCTAAGAGGGATGCACTGT-3’ and 5’-AGACGCATTGACGGATCTTGC-3’ and *Eig73Ee*: 5’-CTAACTGTGGTCTGCTTAGTGG-3’ and 5’-CAACGCTTTCTCAATTACCTCCA-3’.

### Western-blot

Salivary glands from 10 staged larvae were dissected, homogenized with micropestle in Lämmli buffer supplemented with cOmplete (Roche) and DTT. Thereafter, samples were incubated for 10 min at 80°C. 10 µl of lysate was loaded onto a 6% SDS-gel. Proteins were transferred to 0.45 µm nitrocellulose membrane (Biorad: #1620115) and subsequently incubated with PNG-Peroxidase (Sigma-Aldrich #L7759,1:1000) and anti-tubulin (1:5000, AbCam #184970) for 1h at RT. Tubulin was detected with HRP conjugated anti-mouse incubated for 30 min at RT. Bands were detected with ChemiDoc (Biorad) and analyzed for intensity with ImageLab (version 5.2.1). The intensity of 150 kDa bands were normalized to Tubulin. Unpaired t-test was then applied. Coomassie staining was performed according to the protocol (Sigma-Aldrich #B2025)

## Results

### A fibrosis model in *Drosophila* salivary glands

We had previously shown that expression of a dominant-active version of Ras (*Ras*^*V12*^) driven by Beadex-Gal4 (referred here as *Ras*^*V12*^) in salivary glands (SG, [29, 57]) leads to an increase in signal intensity, redistribution and disruption of collagen IV in the BM [46]. Here we initially analyzed the expression of SPARC, which is known to directly interact with collagen IV and found that – similar to collagen IV – its signal intensity increased in *Ras*^*V12*^ SGs (Fig. 1A; quantified in Fig. S1A). The distribution of SPARC and three additional ECM components (Laminin (Lam), Nidogen (Ndg) and Perlecan (Pcan)) was analyzed using specific antibodies without detergent on optical sections close to the surface of SGs where the BM is located (Fig. 1A extracellular), and in the presence of detergent on deeper sections (Fig. 1A, Extra/intracellular). Under both conditions, the regular hexagonal pattern of all BM components in wild type SGs was replaced by a more irregular distribution in *Ras*^*V12*^-expressing glands. All four proteins displayed an increased signal intensity compared to wild-type controls (Fig. 1A). To exclude differences in accessibility to antibodies as an explanation for the difference in signal intensity, we used GFP-tagged collagen IV both for quantification and to measure BM width. This confirmed that compared to control glands, the BM from *Ras*^*V12*^ SGs contained more collagen IV and had also grown thicker (Fig. 1B, C). We conclude that the BM in *Ras*^*V12*^-glands had become disorganized and more extensive, reminiscent of the fibrotic phenotype, which had been observed in *Drosophila* fat bodies [31] and of mammalian fibrotic lesions. Of note, the fibrotic phenotype was most pronounced in the distal parts of SGs similar to other dysplastic features in the same region.

**Fig 1:**
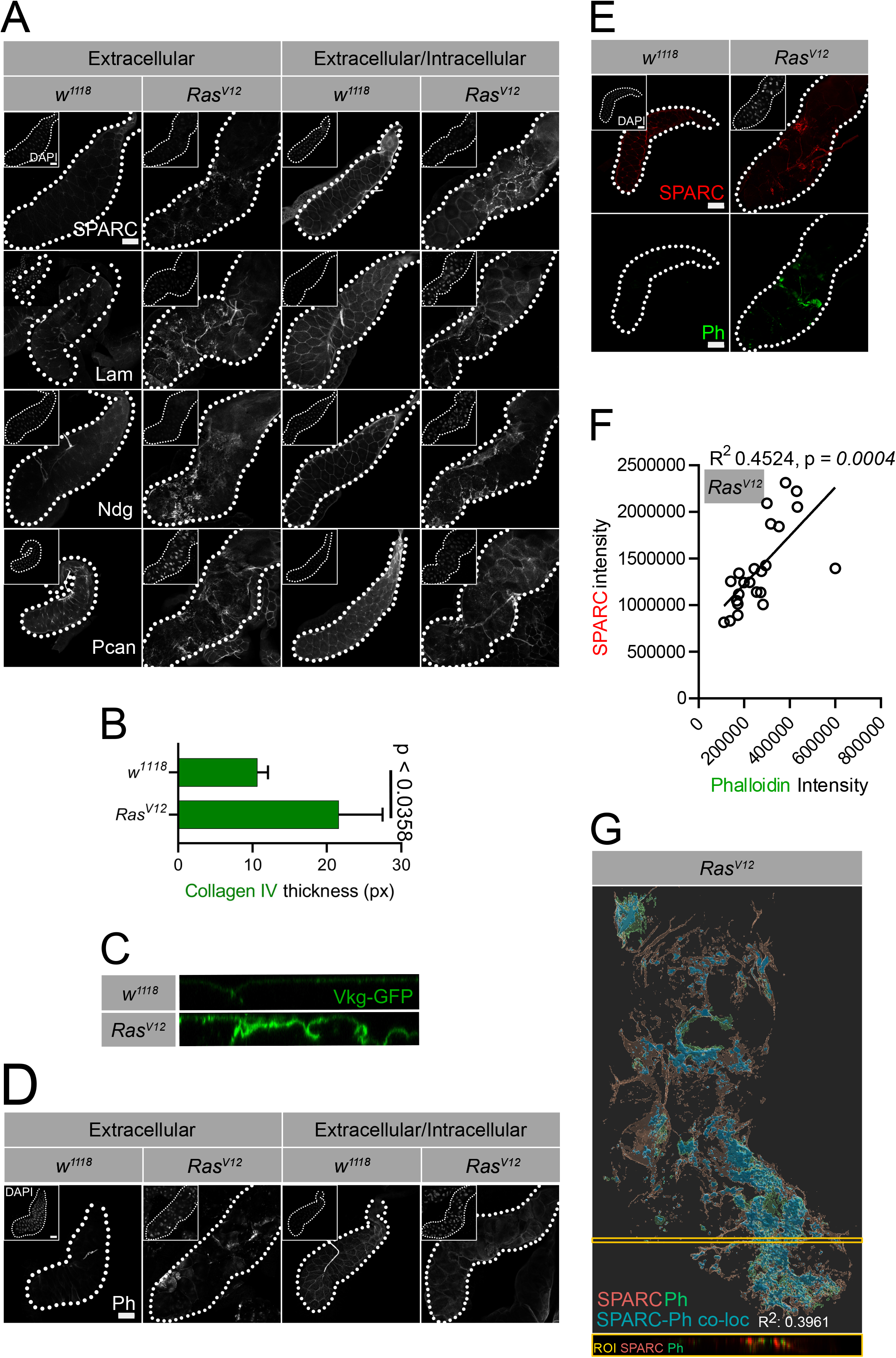
Dysplasia promotes formation of fibrosis and the loss of epithelial organization. (A) PBS and PBST-treated wild type and *Ras*^*V12*^-expressing glands were stained with antibodies against SPARC, Lam, Ndg, Pcan and Actin. (B-C) Quantification of Collagen IV (Vkg-GFP) thickness in *w*^*1118*^ and *Ras*^*V12*^ glands shows an increase in BM thickness (p < 0.0358). (D) Extracellular F-Actin accumulates on the surface of the *Ras*^*V12*^ glands (detected by Phalloidin, Ph). (E-G) The intensity of SPARC and Actin co-vary (E and F; R^2^ 0,4524) and (G) co-localize (R^2^ 0,3961). A Region Of Interest (ROI); is indicated in yellow) and displayed as an unprocessed z-stack showing moderate co-localization of SPARC (Red) and Phalloidin (Green). The scale bar (A, D-E) represents 100 µm.

Although initially used as a control to distinguish between normal and dysplastic SGs, F-actin staining with phalloidin largely mimicked the disorganized distribution of the BM components and was also detected in *Ras*^*V12*^ SGs in the absence of detergent on surface sections, indicating that some F-actin may have obtained access to the extracellular compartment (Fig. 1D quantified in Fig. S1B). This was further confirmed by showing significant co-variation between F-actin and SPARC on the surface of whole glands (Fig. 1E, F). Additionally, by analyzing Z-scanned SGs, we observed a moderate SPARC-phalloidin co-localization (Fig. 1G).

Thus, we find that in the distal part of *Ras*^*V12*^ SGs, BM components are redistributed in concert leading to a fibrotic phenotype including F-actin, part of which may leak out from SG cells and associate with the BM.

### The immune system targets dysplastic salivary glands

To assess the influence of hemostasis on fibrosis, we first measured the activity of *Drosophila* transglutaminase, which is a key clotting factor and has been shown to target intruders such as bacteria and insect pathogenic nematodes [26]. Mammalian TGs are known to be activated in fibrotic lesions in several organs [13, 14, 58, 59]. Activity of the single *Drosophila* TG was detected with an antibody against the covalent ε-(ϒ-L-glutamyl)-L-lysine links that are created by TG and, to focus on extracellular TG, we used non-permeabilized SGs and surface sections (Fig. 2). Compared to wild type glands, significantly higher TG activity was detected on *Ras*^*V12*^ SGs, indicating that - similar to microbial intruders - dysplastic glands are targeted by TG (Fig. 2A, B). In contrast to wild type, where it was faint, TG activity in *Ras*^*V12*^ SGs was found to co-vary (Fig. 2C) and moderately co-localize with Phalloidin intensity (Fig. 2D). In addition to TG, Tiggrin, which is also part of *Drosophila* hemolymph clots [60] was recruited to *Ras*^*V12*^ SGs further confirming that hemostasis is activated against dysplastic SGs (Fig. S1C,D).

**Fig 2:**
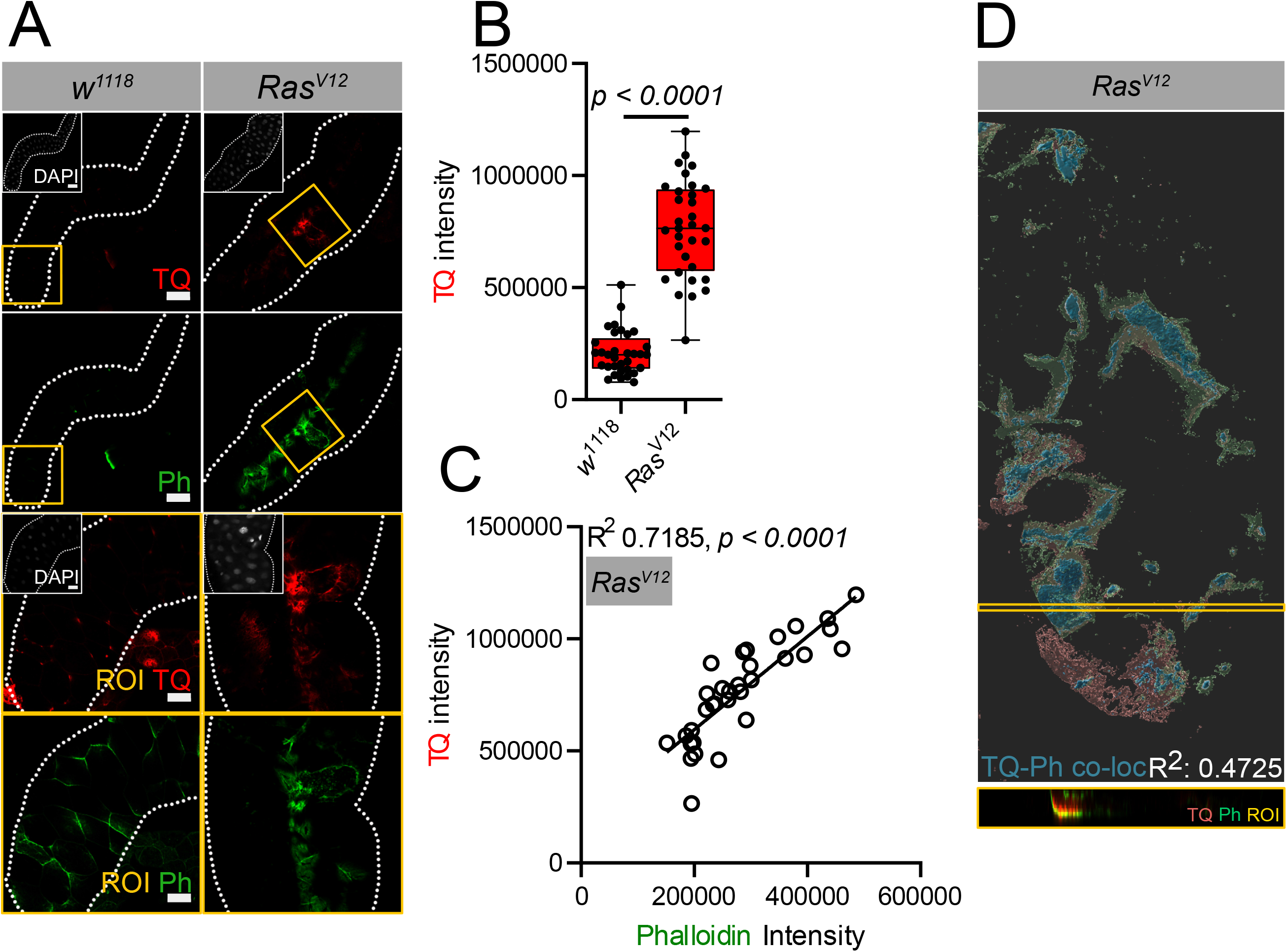
Activation of Transglutaminase activity in *Ras*^*V12*^-expressing glands. (A-B) Higher Transglutaminase activity (TQ in Red) was detected on *Ras*^*V12*^-expressing SGs in comparison to *w*^*1118*^ SGs (quantified in B) Upper ROI insert (A: yellow) are magnified in the lower panel. (C) TG (Red) activity co-varies with Phalloidin (Green, p < 0.0001). (D) Co-localization analysis showed a moderate overlap between TQ and Phalloidin (Turquoise, R^2^ 0.4725). ROI Inset (D: yellow) displays an unprocessed z-stack of TQ (Red) and Phalloidin (Green) The scale bars in (A) correspond to: 100 µm (two upper panels) and 50 µm (ROI).

Ultimately, both coagulation of hemolymph, the formation of nodules and the encapsulation of larger objects result in the recruitment and activation of PPO. During coagulation, this follows TG–activation [22]. To test whether PPOs are also recruited to dysplastic Ras ^V12^- glands, we used antibodies specific for PPO1 and PPO2, the two PPOs expressed by crystal cells and found that both of them were detected on *Ras*^*V12*^–expressing glands (Fig. S1E-G). Further analysis showed that PPO1 intensity and Phalloidin staining co-varied, whereas PPO2 did not (Fig. S1H,I). In summary, these data show that beyond induction of antimicrobial peptides and hemocyte recruitment [46], *Ras* ^*V12*^-SGs display local activation of the hemostatic system. This includes structural clot components such as Tiggrin as well as enzymatic activities such as transglutaminase, followed by recruitment of phenoloxidases.

### Defective secretion in the dysplastic gland

Having observed fibrosis in dysplastic glands prompted us to investigate the integrity and secretory potential of SGs. As indicated by [39], in addition to a disrupted BM, a second precondition for activation of an inflammatory reaction against aberrant tissue is a loss of apico-basal polarity. To test whether SG epithelial cells had undergone similar changes, we assessed the distribution of ß-integrin, F-actin and Dlg in permeabilized SGs. ß-integrin, one of the receptors for BM components, shows an even surface expression and enrichment at the basal side in wild type SG cells. This pattern had been lost upon *Ras*^*V12*^ expression, together with a complete loss of the SG lumen detected by Phalloidin staining (Fig. 3A). Similarly, the regular hexagonal pattern detected with Dics-large (Dlg)-specific antibodies was replaced by a weaker irregular pattern in *Ras*^*V12*^ SGs. To test the secretory potential of *Ras*^*V12*^ SGs we assessed the secretion of two known salivary gland secretory proteins, Sgs3 (salivary glands secreted protein 3) and Eig71Ee [61, 62]. We first investigated the intensity levels of Sgs3 using a GFP-tagged version under its endogenous promoter. In comparison to control SGs, the dysplastic glands displayed lower Sgs3 levels, in line with the absence of a lumen (Fig. 3B, C). Western-blotting and Coomassie staining revealed lower levels of Eig71Ee in *Ras*^*V12*^ glands, in comparison to control glands (Fig. 3D, S2A). Altogether, dysplastic glands displayed a loss of epithelial character and produced lower amounts of two main secretory proteins. Nevertheless, most proteins were still shared with wild type glands and no general degradation was observed (Fig. S2A).

**Fig 3:**
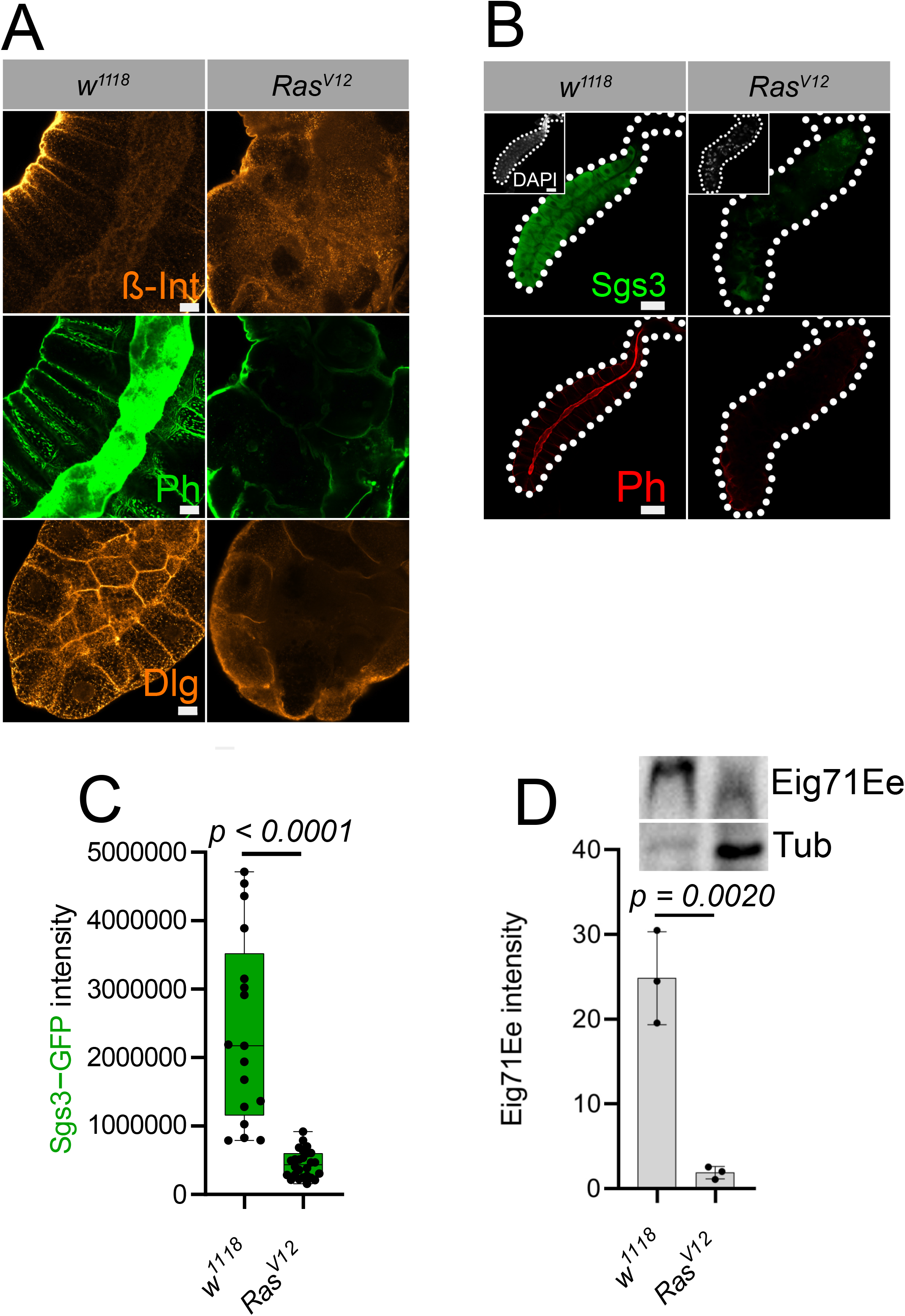
*Ras*^*V12*^ glands display a partial loss of epithelial character and loss of secretion. (A) In *Ras*^*V12*^ SGs, ß-integrin is lost from the basolateral site, F-Actin (Phalloidin) cuboidal distribution is lost and Dlg is absent from the apical site, in comparison to *w*^*1118*^ glands. (B-C) Sgs3 intensity is reduced in *Ras*^*V12*^ glands detected by Sgs3-GFP (B, quantified in C). (D) Western-blot analysis shows a higher amount of Eig71Ee in *w*^*11118*^ in comparison to *Ras*^*V12*^ SGs (p = 0.002, N=3, D). Scale bar represent in (A) 50 µm and in (B) 100 µm.

### An antimicrobial peptide reverts BM dysplasia, loss of epithelial character and secretion

We had previously found that Drs is expressed most strongly in the proximal region of *Ras*^*V12*^-glands, a region which displayed fewer signs of fibrosis and looked more similar to wild type SGs. We had also managed to revert collagen IV deregulation by expressing Drs across whole glands including the strongly affected distal part [46], while expression of control GFP [29] and membrane-bound RFP [46] under the same UAS control had no such effects. Therefore, we wondered whether the fibrotic distribution of other BM components was be rescued upon forced Drs expression across whole glands. Indeed, in *Drs-*overexpressing *(OE)* Ras^V12^ SGs, SPARC, Laminin, Nidogen, Perlecan and Actin displayed a more regular localization, closer to the wild type situation in surface sections (Fig. 4A, B and S2B). This was not observed upon RFP overexpression excluding a dilution effect due to the presence of dual UAS targets (Fig. S2D). In line with the more regular distribution of BM components, BM thickness in *Drs-*overexpressing *Ras* ^*V12*^ SGs was also reduced to wild type levels (Fig. 4C, D). Moreover, transglutaminase activity and PPO1 binding (Fig. 4E-G) decreased, although not completely, when compared to *Ras* ^*V12*^-expressing SGs, whereas PPO2 (Fig. S2C) did not revert to wild type levels. At the same time, staining with Phalloidin confirmed that F-Actin had been reduced to almost wild type levels (Fig. 4B).

**Fig 4:**
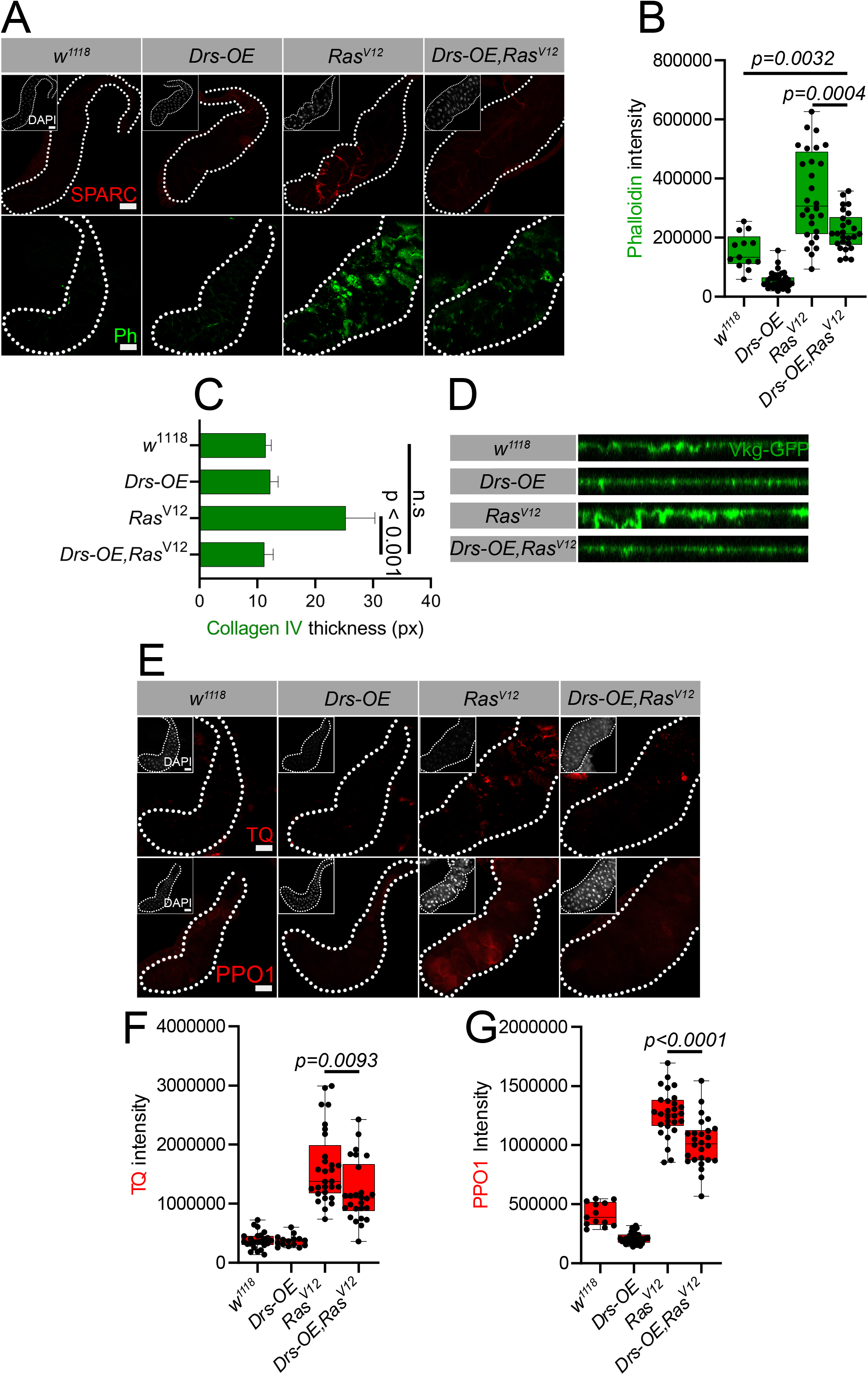
Drosomycin expression prevents fibrosis in *Ras*^*V12*^ glands. (A) Drosomycin overexpression in *Ras*^*V12*^ glands (*Drs-OE,Ras*^*V12*^) leads to reduced expression of SPARC and F-Actin (Ph, quantified in B: p = 0.0004) compared to *Ras*^*V12*^ SGs. (C-D) Collagen IV thickness was restored to control levels in *Drs-OE,Ras*^*V12*^ glands, in comparison to *Ras*^*V12*^ SGs (p < 0.001). (E-F) TG activity (TQ, quantified in F: p = 0.0093) and PPO1 (E, quantified in G: p > 0.0001), in comparison to *Ras*^*V12*^ SGs was also reduced upon co-expression of Drs. Scale bars in (A and D) represent 100 µm.

To examine the secretory activity in normal, *Ras*^*V12*^-expressing and *Drs-OE* SGs we compared SG histology and the production of Sgs3 and Eig71Ee. In line with the renewal of a more normal BM morphology and the restoration of the epithelial character, as assayed by integrin, Dlg and Phalloidin staining (Fig. 5A), the lumen of *Drs-OE, Ras*^*V12*^ SGs was also partially restored allowing production of both Sgs3 (Fig. 5B, C and E) and Eig71Ee (Fig. 5D, E), in comparison to *Ras*^*V12*^ glands (Fig. 5B). *Sgs3* levels in *Ras*^*V12*^ SGs were not restored when RFP was co-expressed as a negative control (Fig. S2F). Both Western-blot (Fig. 5D) confirmed higher levels of Eig71Ee in *Drs-OE, Ras*^*V12*^ glands.

**Fig 5:**
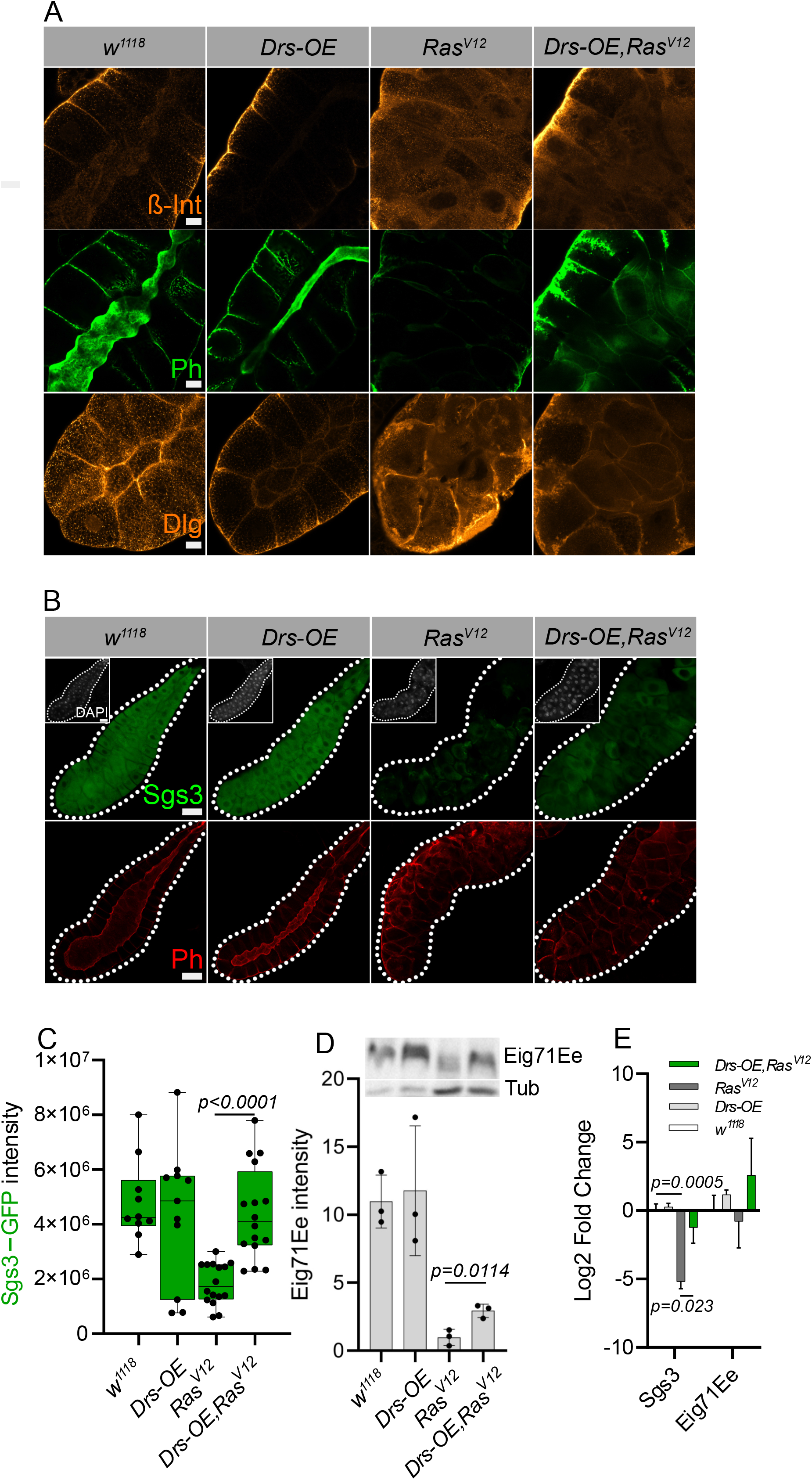
The epithelial character and production of secreted proteins is restored upon co-expressing Drs in *Ras*^*V12*^ glands. (A) Staining of control (w1118; and Drs-overexpressing, (*Drs-OE)*), *Ras*^*V12*^-expressing as well as Drs co-expressing glands (*Drs-OE, Ras*^*V12*^) with Integrin-specific antibodies, Phalloidin and Dlg shows a partial reversion to an epithelial phenotype. (B) The production of Sgs3 (Sgs3-GFP) is restored in *Drs-OE, Ras*^*V12*^ glands compared to *Ras*^*V12*^ glands (quantified in C: p < 0.0001) in line with the restoration of the SG lumen (detected with Phalloidin). (D) Western-blot analysis shows increased production of Eig71Ee (gp150, N=3), in *Drs-OE,Ras*^*V12*^ glands (D: p < 0.0114) (gp150, N=3). (E) qRT-PCR analysis of *Sgs3* and *Eig71Ee* show increased expression in *Drs-OE,Ras*^*V12*^ glands in comparison to *Ras*^*V12*^. Scale bars represent 20 µm (A) and 100 µm (B).

### Drs localization in fibrotic lesions

To detect whether Drs is properly located to interfere with the formation of fibrotic lesions in *Ras*^*V12*^ SGs, we used an HA-tagged version (Drs-HA) to analyze surface sections (Fig. 6). Since it had previously been shown that AMPs have the potential to bind Actin [63] and based on our own results (see above), we used Phalloidin as a proxy to identify fibrotic lesions. Signals for both Phalloidin and Drs-HA were detected in *Ras*^*V12*^ SGs and a weak signal upon expression of Drs-HA alone in a wild type background (Fig. 6A). In both cases, Drs-HA overlapped with the Phalloidin signal, indicating partial co-localization with F-Actin, which was confirmed in Z-sections (Fig. 6B). Taken together, the distribution of Drs in *Ras*^*V12*^ SGs is compatible with its inhibitory effect on fibrotic lesions.

**Fig 6:**
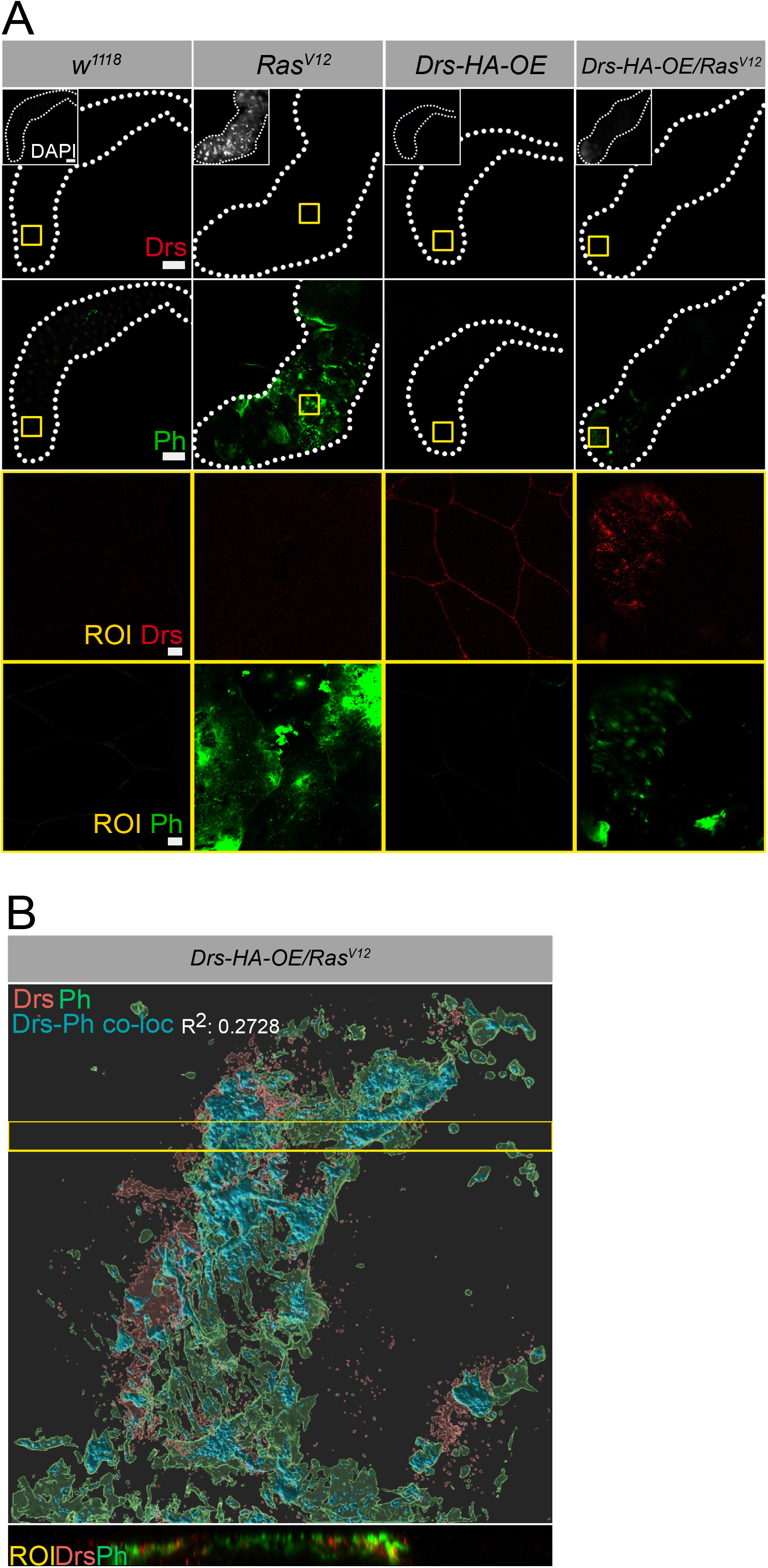
Drs co-localizes with F-Actin in fibrotic lesions. (A) HA-tagged Drs (Drs-HA) expressed in wild type and *Ras*^*V12*^*-*expressing SGs was detected using HA-specific antibodies as well as Phalloidin. Then upper ROI inserts (A: yellow) are magnified in the lower panels. (B) Partial overlap between the signals was confirmed using confocal microscopy including Z-stacks. ROI Inset (B: yellow) displays an unprocessed Z-stack of Drs-HA (Red) and Phalloidin (Green)

## Discussion

We show that expression of a dominant active form of an oncogene induces a fibrotic phenotype at the basement membrane of *Drosophila* salivary glands, a highly active secretory organ (summarized in Fig. 7). In contrast to other *Drosophila* fibrosis models, the redistribution of BM components, which are primarily produced in the fat body ([38], see also supplementary table S2) and deposited onto internal organs, is most likely a secondary consequence of SG dysplasia. Although we observe an increase in signal intensity for all BM components in *Ras*^*V12*^ SGs, this appears not to be a consequence of transcriptional activation, since the BM components are only poorly if at all induced in the fat body of *Ras*^*V12*^ larvae (see supplementary table S2). As a proxy for BM components, SPARC is found associated with extracellular F-Actin, which appears to be released from dysplastic cells either SG cells or rupturing crystal cells [22]. When analyzing Transglutaminase activity as a proxy for hemostasis, we found increased levels in *Ras*^*V12*^ SGs. Similar to what happens during wound closure in flies, TG activation was followed by recruitment of Phenoloxidase, although in this case only the PPO1 signal co-varied with Phalloidin staining of F-Actin. Taken together we detect fibrotic lesions, which contain BM components, TG activity, PPO1 and extracellular F-Actin. These findings are in line with the proposed model for fibrotic lesions as unabated wound healing. Although we were initially motivated by a possible hemostatic/immune function of TG, the single *Drosophila* TG may play a slightly different role during fibrosis similar to mammalian tissue transglutaminase (TG2), which has been shown to aggravate fibrosis in several organs due to its crosslinking activity [13, 14, 58, 59].

**Fig 7:**
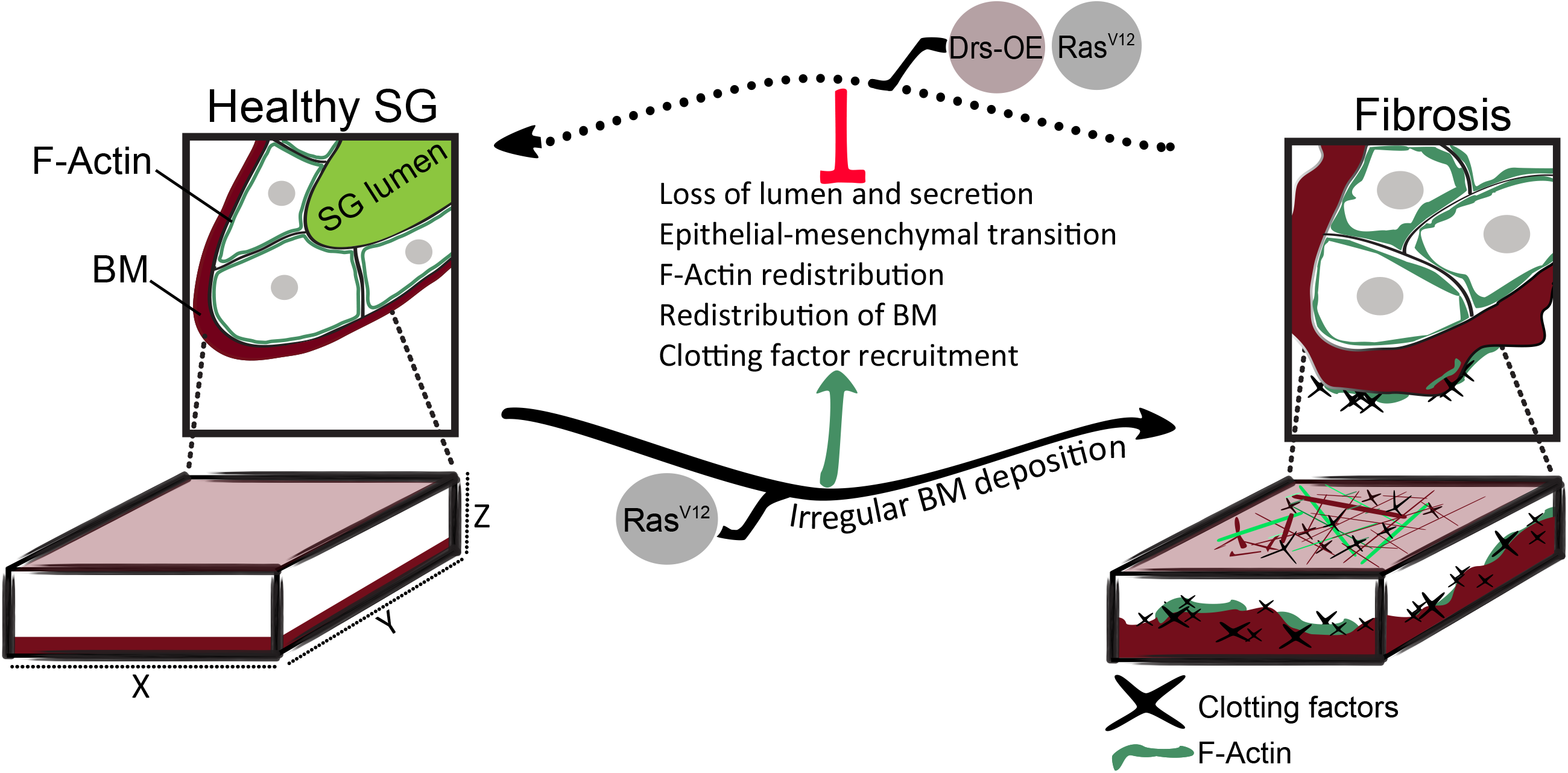
Fibrosis in salivary glands and its reversal through Drosomycin expression. The basal membrane (BM, represented below as in confocal stacks), of *Ras*^*V12*-^expressing salivary glands (SGs) shows several features of fibrotic lesions, which are substantially reversed upon overexpression of the antimicrobial peptide Drosomycin. Fibrotic lesions incorporate clotting factors from the hemolymph-facing (basal) layer and F-actin from underlying SG cells (see text for further details).

SG fibrosis affects the normal secretory function of SGs as shown here by (1) a loss of the epithelial character of SG cells (2) the collapse of the SG lumen and (3) reduced production of two secretory proteins (Sgs3 and Eig71Ee).

Supporting our previous observations that single expression of the AMP Drosomycin rescues several features of the dysplastic phenotype [46], we find that upon Drs overexpression, the formation of fibrotic lesions at the BM is substantially reduced and tissue integrity is largely restored including a partial reversion to an epithelial character of SG cells and restoration of secretory activity (summarized in Fig. 7). This is paralleled by a reduction in hemostatic/immune activation including (1) fewer TG crosslinks and (2) a reduction in recruitment of PPO1, but not PPO2. These findings support the notion that early hemostatic components are recruited towards fibrotic foci, similar to wounds were PPO1 and TG activity have been shown to precede PPO2, which is subsequently released by rupture of crystal cells [64, 22].

Finally, we observe that Drs co-localizes with F-Actin in fibrotic lesions similar to the mammalian AMP LL-37 which has previously been shown to interact with F-Actin [65, 63] with different effects including protection from microbial degradation [66], but also inhibition of antimicrobial activity [67]. Due to high background staining, we were unable to determine whether Drs is also secreted apically into the SG lumen. Therefore, we do not know whether Drs equally influences apical and basal secretion of SG cells. For the basal localization, we propose a scenario, where secreted Drs reduces the formation of fibrotic plaques by interfering with the aggregation of BM components. These may include particulate Actin (F-Actin), which, once released from cells, is known to have deleterious effects, for example during sepsis [68], by acting as a procoagulant [69], and through inhibition of macrophage defenses [70]. These effects are counteracted by Actin-scavenging systems [69], which include soluble Gelsolin [71], Vitamin D-binding protein [72] and – as proposed here - Drs. In line with potential role as damage-associated molecular patterns [73-75] cytosolic proteins have been shown to activate immunity in flies [76, 77]. Irrespective of its target in fibrotic foci, the anti-fibrotic effects of Drs are of clinical interest and may include other AMPs such as the bee AMP Melittin [78-80], which restores the epithelial character [81] similar to what we observe. Taken together, the data presented here support the concept that dysplasia of tumor cells supports the formation of fibrotic aggregates [10, 6]. The recruitment of both TG and PO, which are known players in hemolymph coagulation lends further support to the concept that tumors induce a chronic procoagulatory state [82, 83] and adds the notion that AMPs have the potential to revert this phenotype.

## Acknowledgments

We thank Stina Höglund, Chris Molenaar and the Imaging facility at Stockholm University for support with all aspects of microscopy. We also thank the anymous reviewers for their constructive feedback, Roger Karlsson for his critical input and the late Thomas Stössel for drawing our attention to actin-scavenging systems.

## Statement of Ethics

No approvement of studies involving animals was required.

## Conflict of Interest Statement

The authors declare no conflict of interest.

## Funding Sources

This work was supported by the Swedish Research Council (VR-2010-5988 and VR 2016-04077) and the Swedish Cancer Foundation (CAN 2010/553 and CAN 2013/546) to UT and the Eric Philipp Sörensen Stiftelse to SB.

## Author Contributions

experiment design: DK, SB and UT; experimental work: DK, CK and SB; raising funding: SB and UT; manuscript drafting: DK, SB and UT.

## Notes

### Competing Interest Statement

The authors have declared no competing interest.

