## Supplemental figures and tables for "Anti-fibrotic activity of an antimicrobial peptide in a *Drosophila* model"

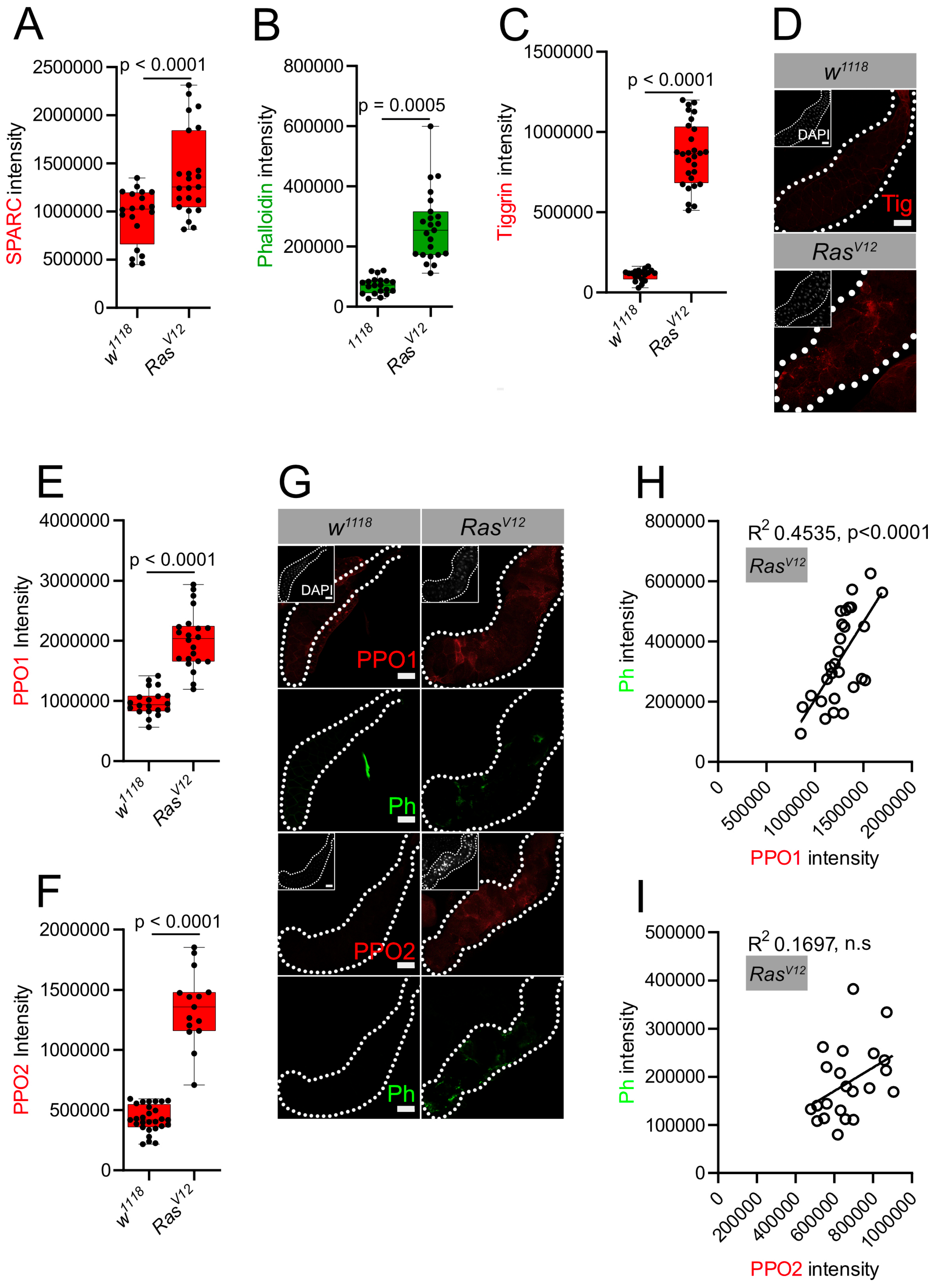


**Supplementary Figure S1**. Fibrotic formation in *Ras*$^{V12}$ salivary glands. (A-C) FI (fluorescence intensity) quantification of *w*$^{1118}$ and *Ras*$^{V12}$ salivary glands stained for SPARC ($p < 0.0001$), Phalloidin ($p = 0.0005$) and Tiggrin ($p < 0.0001$). (D) Representative images were stained for Tiggrin. (E-F) FI quantification of *w*$^{1118}$ and *Ras*$^{V12}$ salivary glands stained for PPO1 ($p < 0.0001$) and PPO2 ($p < 0.0001$).(G) Representative images were stained for PPO1, Phalloidin and PPO2 (H-I) Co-variation analysis of PPO1 and Phalloidin (ph; $p < 0.0001$), and (G) PPO2 with Ph (n.s). Scale bar represents 100 μm (D and G)

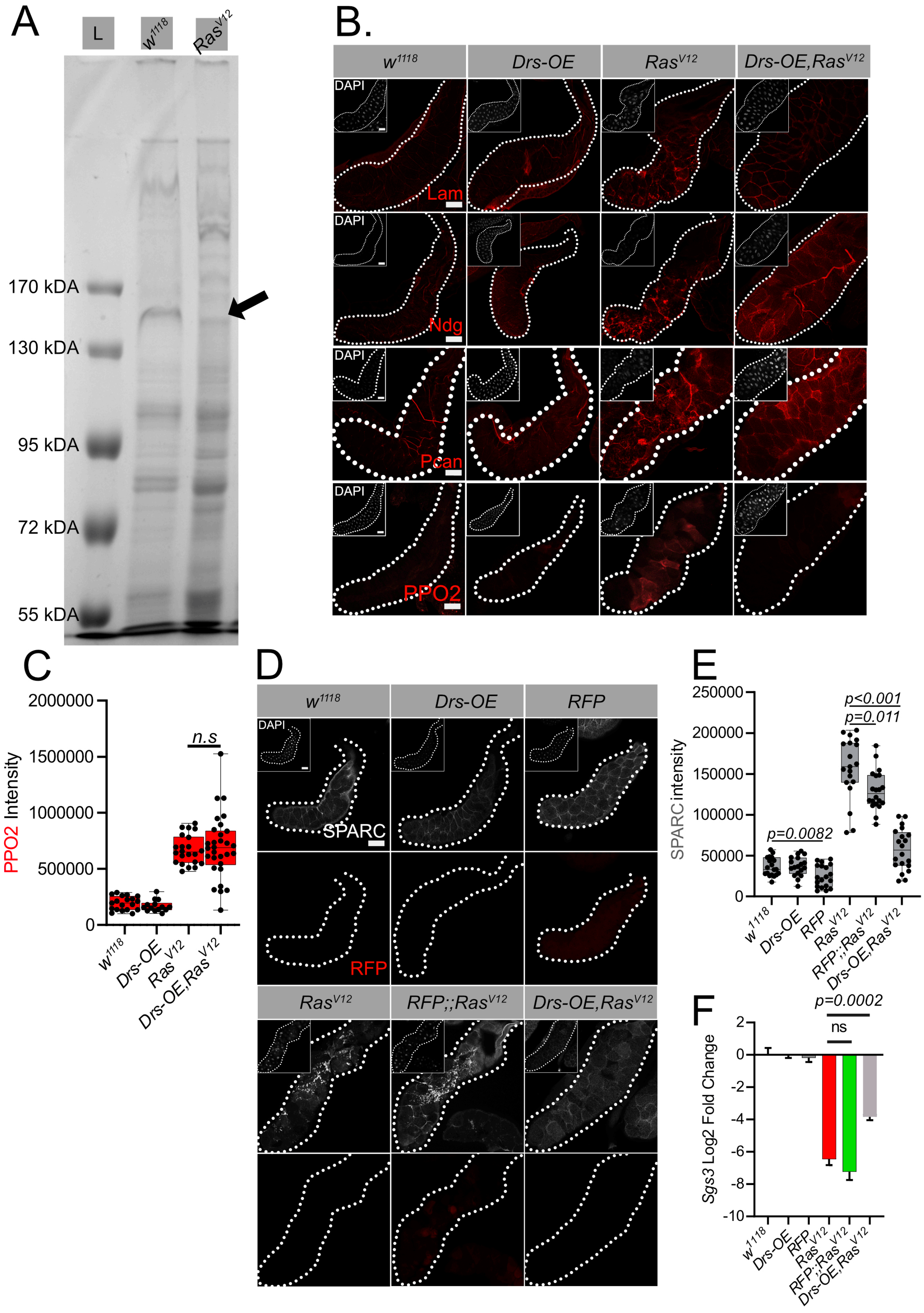


Supplementary Figure S2: (A) Coomassie staining shows a reduction in the intensity of the 150kDA band (gp150, indicated by the arrow; N=3). (B) *Drs-OE* in *Ras$^{V12}$* glands restores distribution of Lam, Ndg, Pcan. In contrast, PPO2 is still recruited to a similar extent (quantified in C: n.s). (D) The effect of *Drs* on SPARC accumulation is not dependent on the presence of dual UAS targets, since SGs expressing *RFP* under UAS control (*RFP*;;*Ras$^{V12}$*) retained the fibrotic phenotype (quantified in E, note that upon expression of *RFP*, a slight reduction in SPARC intensity is observed in both wild type and *Ras$^{V12}$* lines). (F) qPCR analysis shows Sgs3 levels are affected by overexpression of *Drs* and not *RFP* in *Ras$^{V12}$* glands. Scale bar represents 100 μm (B and D).

**Supplementary table S1**. List of all crosses corresponding to the respective figures.

**Figure 1.**
A-E
♀ *w1118,Beadex-Gal4 (Bx)* > ♂ $w^{1118}$
♀ *Bx* > ♂ $w^{1118}$;;*UAS-*$Ras^{V12}$ ($Ras^{V12}$)

F-G
♀ *Bx* >♂ $Ras^{V12}$

**Figure 2.**
A-B
♀ *Bx* > ♂ $w^{1118}$
♀ *Bx* > ♂ $Ras^{V12}$

C-D
♀ *Bx* > ♂ $Ras^{V12}$

**Figure 3.**
A-D
♀ *Bx* > ♂ $w^{1118}$
♀ *Bx* > ♂ $Ras^{V12}$

**Figure 4.**
A-G
♀ *Bx* > ♂ $w^{1118}$
♀ *Bx* > ♂ $w^{1118}$;;*UAS-Drs-OE* (*Drs-OE*)
♀ *Bx* > ♂ $Ras^{V12}$
♀ *Bx* > ♂ $w^{1118}$;;*UAS-Drs-OE,UAS-*$Ras^{V12}$ (*Drs-OE,*$Ras^{V12}$)

**Figure 5.**
A-E
♀ *Bx* > ♂ $w^{1118}$
♀ *Bx* > ♂ *Drs-OE*
♀ *Bx* > ♂ $Ras^{V12}$
♀ *Bx* > ♂ *Drs-OE,*$Ras^{V12}$

**Figure 6.**
A
♀ *Bx* > ♂ $w^{1118}$
♀ *Bx* > ♂ $Ras^{V12}$
♀ *Bx;;UAS-Drs-HA* > ♂ $w^{1118}$
♀ *Bx;;UAS-Drs-HA* > ♂ $Ras^{V12}$

B
♀ *Bx;;UAS-Drs-HA* > ♂ $Ras^{V12}$

**Figure S1.**

A-G

♀*Bx* > ♂$w^{1118}$

♀*Bx* > ♂$Ras^{V12}$

H-I

♀*Bx* > ♂$Ras^{V12}$

**Figure S2.**

A

♀*Bx* > ♂$w^{1118}$

♀*Bx* > ♂$Ras^{V12}$

B-C

♀*Bx* > ♂$w^{1118}$

♀*Bx* > ♂*Drs-OE*

♀*Bx* > ♂$Ras^{V12}$

♀*Bx* > ♂*Drs-OE,*$Ras^{V12}$

D-F

♀*Bx* > ♂$w^{1118}$

♀*Bx* > ♂*Drs-OE*

♀*Bx,UAS-mCD8::RFP* > ♂$w^{1118}$

♀*Bx* > ♂$Ras^{V12}$

♀*Bx,UAS-mCD8::RFP* > ♂$Ras^{V12}$

♀*Bx* > ♂*Drs-OE,*$Ras^{V12}$

Supplementary table S2. ECM expression pattern. *SPARC, Lam A, Ndg, Pcan* and *Vkg* expression in fat body (FB) and salivary glands (SG) of wandering larvae, and in wild type compared to $Ras^{V12}$-expressing FBs and SGs (96 h after egg deposition, transcript counts and Log2 of the relative differences are indicated), Data derived from FlyAtlas and our previously published work.

| Genes | FB FlyAtlas [L3 wandering: Linear] | SG FlyAtlas [L3 wandering: Linear] | FB seq [w1118 vs RasV12] Log2 | SG seq [w1118 vs RasV12] Log2 |
|---|---|---|---|---|
| Sparc (BM40) | 500 | 20 | [814 vs 755,6]-0,1 | N/A |
| Lam A | 53 | 1 | [209,3 vs 236,4]0,17 | N/A |
| Ndg | 12 | 0 | [1,48 vs 12,54]3,07 | [0,06 vs 0,24]4,07 |
| Trol (Pcan) | 169 | 2 | [572 vs 397] -0,54 | [64 vs 24]0,42 |
| Vkg | 83 | 1 | [302 vs 152] -0,99 | N/A |
